# Concurrent Multisensory Integration and Segregation with Complementary Congruent and Opposite Neurons

**DOI:** 10.1101/471490

**Authors:** Wen-Hao Zhang, He Wang, Aihua Chen, Yong Gu, Tai Sing Lee, K. Y. Michael Wong, Si Wu

**Affiliations:** Department of Physics, Hong Kong University of Science and Technology, Hong Kong.; School of Electronics Engineering and Computer Science, IDG/McGovern Institute for Brain Research, Peking University, Beijing 100871, China.; The Center of the Neural Basis of Cognition, Carnegie Mellon University, Pittsburgh, PA 15213.; Laboratory of Brain Functional Genomics, Primate Research Center, East China Normal University, Shanghai, China.; Institute of Neuroscience, Chinese Academy of Sciences, Shanghai, China.

**Keywords:** Opposite neuron, Multisensory integration, Concurrent integration and segregation, Decentralized architecture, Continuous attractor neural network

## Abstract

Our brain perceives the world by exploiting multiple sensory modalities to extract information about various aspects of external stimuli. If these sensory cues are from the same stimulus of interest, they should be integrated to improve perception; otherwise, they should be segregated to distinguish different stimuli. In reality, however, the brain faces the challenge of recognizing stimuli without knowing in advance whether sensory cues come from the same or different stimuli. To address this challenge and to recognize stimuli rapidly, we argue that the brain should carry out multisensory integration and segregation concurrently with complementary neuron groups. Studying an example of inferring heading-direction via visual and vestibular cues, we develop a concurrent multisensory processing neural model which consists of two reciprocally connected modules, the dorsal medial superior temporal area (MSTd) and the ventral intraparietal area (VIP), and that at each module, there exists two distinguishing groups of neurons, congruent and opposite neurons. Specifically, congruent neurons implement cue integration, while opposite neurons compute the cue disparity, both optimally as described by Bayesian inference. The two groups of neurons provide complementary information which enables the neural system to assess the validity of cue integration and, if necessary, to recover the lost information associated with individual cues without re-gathering new inputs. Through this process, the brain achieves rapid stimulus perception if the cues come from the same stimulus of interest, and differentiates and recognizes stimuli based on individual cues with little time delay if the cues come from different stimuli of interest. Our study unveils the indispensable role of opposite neurons in multisensory processing and sheds light on our understanding of how the brain achieves multisensory processing efficiently and rapidly.

**Significance Statement:** Our brain perceives the world by exploiting multiple sensory cues. These cues need to be integrated to improve perception if they come from the same stimulus and otherwise be segregated. To address the challenge of recognizing whether sensory cues come from the same or different stimuli that are unknown in advance, we propose that the brain should carry out multisensory integration and segregation concurrently with two different neuron groups. Specifically, congruent neurons implement cue integration, while opposite neurons compute the cue disparity, and the interplay between them achieves rapid stimulus recognition without information loss. We apply our model to the example of inferring heading-direction based on visual and vestibular cues and reproduce the experimental data successfully.

## Introduction

To survive as an animal is to face the daily challenge of perceiving and responding fast to a constantly changing world. The brain carries out this task by gathering as much as possible information about external environments via adopting multiple sensory modalities including vision, audition, olfaction, tactile, vestibular perception, etc. These sensory modalities provide different types of information about various aspects of the external world, and serve as complementary cues to improve perception in ambiguous conditions. For instance, while walking, both the visual input (optic flow) and the vestibular signal (body movement) convey useful information about heading-direction, and when integrated together, they give a more reliable estimate of heading-direction than either of the sensory modalities could deliver on its own. Indeed, experimental data has shown that the brain does integrate visual and vestibular cues to infer heading-direction and furthermore the brain does it in an optimal way as predicted by Bayesian inference ^1^. Over the past years, experimental and theoretical studies verified that optimal information integration were found among many sensory modalities, for example, integration of visual and auditory cues for inferring object location ^2^, motion and texture cues for depth perception^3^, visual and proprioceptive cues for hand position ^4^, and visual and haptic cues for object height ^5^.

However, multisensory integration is only a part of multisensory information processing. While it is appropriate to integrate sensory cues from the same stimulus of interest (Fig. 1A left), sensory cues from different stimuli need to be segregated rather than integrated in order to distinguish and recognize individual stimuli (Fig. 1A right). In reality, the brain does not know in advance whether the cues are from the same or different objects. To recognize stimuli rapidly, we argue that the brain should carry out multisensory integration and segregation concurrently: a group of neurons integrates sensory cues, while the other computes the disparity between cues. The interplay between the two groups of neurons determines the final choice of integration versus segregation.

**Figure 1:**
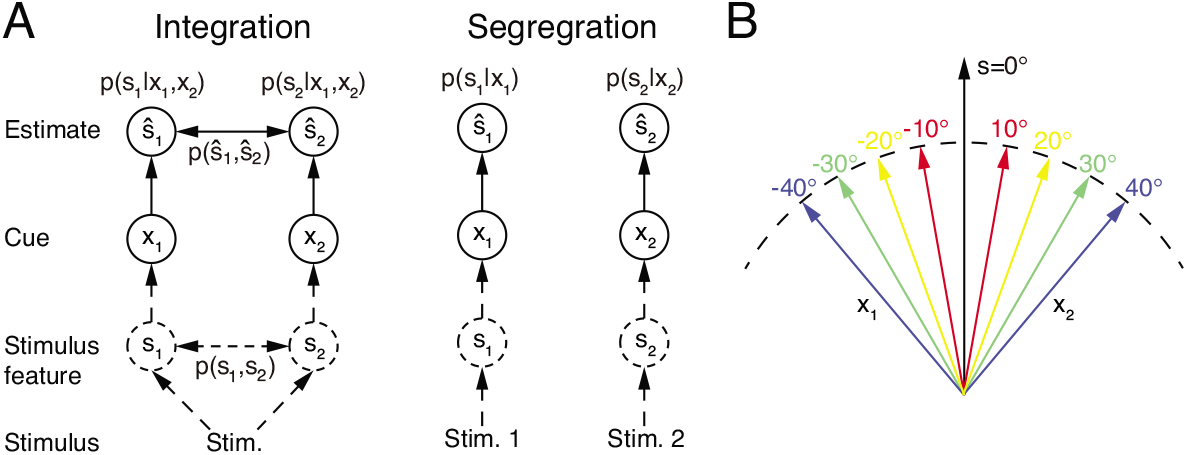
Multisensory integration and segregation. (A) Multisensory integration versus segregation. Two underlying stimulus features *s*_1_ and *s*_2_ independently generate two noisy cues *x*_1_ and *x*_2_, respectively. If the two cues are from the same stimulus, they should be integrated, and in the Bayesian framework, the stimulus estimation is obtained by computing the posterior *p*(*s*_1_|*x*_1_, *x*_2_) (or *p*(*s*_2_ |*x*_1_, *x*_2_)) utilizing the prior knowledge *p*(*s*_1_, *s*_2_) (left). If two cues are from different stimuli, they should be segregated, and the stimulus estimation is obtained by computing the posterior *p*(*s*_1_ |*x*_1_) (or *p*(*s*_2_ |*x*_2_)) using the single cues (right). (B) Information of single cues is lost after integration. The same integrated result *ŝ* = 0° is obtained after integrating two cues of opposite values (θ and -θ) with equal reliability. Therefore, from the integrated result, the values of single cues are unknown.

An accompanying consequence of multisensory integration is, however, that it inevitably incurs information loss of individual cues (Fig. 1, also see SI and Fig. S1). Consider the example of integrating the visual and vestibular cues to infer heading-direction, and suppose that both cues have equal reliability. Given that one cue gives an estimate of *θ* degree and the other an estimate of −*θ* degree, the integrated result is always 0 degree, irrespective to the value of *θ* (Fig. 1B). Once the cues are integrated, the information associated with each individual cue (the value of *θ*) is lost, and the amount of loss information increases with the extent of integration (see SI). Thus, if only multisensory integration is performed, the brain faces a chicken and egg dilemma in stimulus perception: without integrating cues, it may be unable to recognize stimuli reliably in an ambiguous environment; but once cues are integrated, the information from individual cues is lost. Concurrent multisensory integration and segregation is able to disentangle this dilemma.

The information of individual cues can be recovered by using the preserved disparity information if necessary, instead of re-gathering new inputs from the external world. While there are other brain regions processing unisensory information, concurrent multisensory integration and segregation provides an additional way to achieve rapid stimulus perception if the cues come from the same stimulus of interest, and differentiate and recognize stimuli based on individual cues with little time delay if the cues come from different stimuli of interest. This processing scheme is consistent with an experimental finding which showed that the brain can still sense the difference between cues in multisensory integration^6,7^.

What are the neural substrates for implementing concurrent multisensory integration and segregation? Previous studies investigating the integration of visual and vestibular cues to infer heading-direction found that in each of two brain areas, namely, the dorsal medial superior temporal area (MSTd) and the ventral intraparietal area (VIP), there are two types of neurons with comparable number displaying different multisensory behaviors: congruent and opposite cells (Fig. 2)^8,9^. The tuning curves of a congruent cell in response to visual and vestibular cues are similar (Fig. 2A), whereas the tuning curve of an opposite cell in response to a visual cue is shifted by 180 degrees (half of the period) compared to that in response to a vestibular cue (Fig. 2B). Data analysis and modeling studies suggested that congruent neurons are responsible for cue integration^8,10–12^. However, the computational role of opposite neurons remains largely unknown. They do not integrate cues as their responses hardly change when a single cue is replaced by two cues with similar directions. Interestingly, however, their responses vary significantly when the disparity between visual and vestibular cues is enlarged ^13^, indicating that opposite neurons are associated with the disparity information between cues.

**Figure 2:**
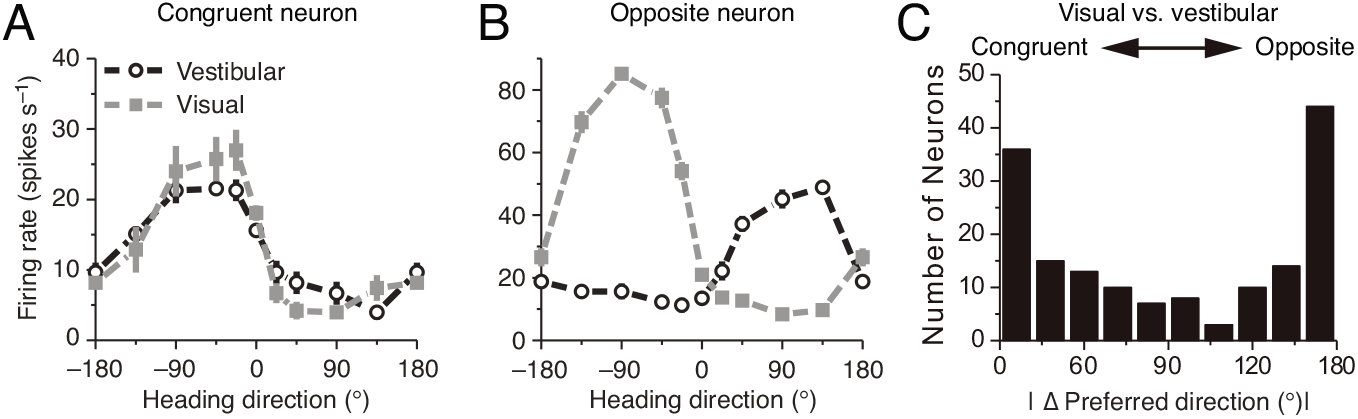
Congruent and opposite neurons in MSTd. Similar results were found in VIP ^18^. (A-B) Tuning curves of a congruent neuron (A) and an opposite neuron (B). The preferred visual and vestibular directions are similar in (A) but are nearly opposite by 180° in (B). (C) The histogram of neurons according to their difference between preferred visual and vestibular directions. Congruent and opposite neurons are comparable in numbers. (A-B) adapted from ref. 8, (C) from ref. 19.

In the present study, we explore whether opposite neurons are responsible for cue segregation in multisensory information processing. Experimental findings showed that many, rather than a single, brain areas exhibit multisensory processing behaviors and that these areas are intensively and reciprocally connected with each other^8,9,14–16^. The architecture of these multisensory areas is consistent with the structure of a decentralized model ^11^, which successfully reproduces almost all known phenomena observed in the multisensory integration experiments^1,17^. Thus we also consider a decentralized multisensory processing model ^11^ in which each local processor receives a direct cue through feedforward inputs from the connected sensory modality and meanwhile, accesses information of other indirect cues via reciprocal connections between processors.

As a working example, we focus on studying the inference of heading-direction based on visual and vestibular cues. The network model consists of interconnected MSTd and VIP modules, where congruent and opposite neurons are widely found^8,9^. Specifically, we propose that congruent neurons in the two brain areas are reciprocally connected with each other in the congruent manner: the closer between the peferred directions of a pair of neurons in their respective brain areas, the stronger their connection is, and this connection profile encodes effectively the prior knowledge about the two cues coming from the same stimulus. On the other hand, opposite neurons in the two brain areas are reciprocally connected in the opposite manner: the further away between the preferred directions of a pair of neurons in their respective brain areas (the maximal difference is 180 degree), the stronger their connection is. Our model reproduces the tuning properties of opposite neurons, and verifies that opposite neurons encode the disparity information between cues. Furthermore, we demonstrate that this disparity information, in coordination with the integration result of congruent neurons, enables the neural system to assess the validity of cue integration and to recover the lost information of individual cues if necessary. Our study sheds light on our understanding of how the brain achieves multisensory information processing efficiently and rapidly.

## Results

### Probabilistic models of multisensory processing

The brain infers stimulus information based on ambiguous sensory cues. We therefore formulate the multisensory processing problem in the framework of probabilistic inference, and as a working example, we focus on studying the inference of heading-direction based on visual and vestibular cues.

#### Probabilistic model of multisensory integration

To begin with, we introduce the probabilistic model of multisensory integration. Suppose two stimulus features {*s_m_* } generate two sensory cues {*x_m_* }, for *m* = 1, 2 (the visual and vestibular cues) respectively (Fig. 1A), and we denote the corresponding likelihood functions as *p*(*x_m_* |*s_m_*). The task of multisensory processing is to infer {*s_m_* } based on {*x_m_* }. *x_m_* is referred to as the direct cue of *s_m_* (e.g., the visual cue to MSTd) and *x_l_* (*l ≠ m*) the indirect cue of *s_m_* (e.g., the vestibular cue to MSTd).

Since heading-direction is a circular variable in the range of (–π, π], we adopt the von Mises, rather than the Gaussian, distribution to carry out the theoretical analysis. In the form of the von Mises distribution, the likelihood function is given by

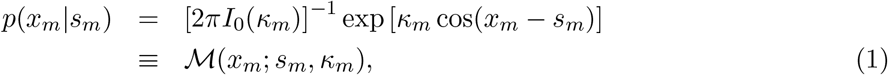

where *I*_0_ (*κ*;) is the modified Bessel function of the first kind and order zero, and acts as the normalization factor. *s_m_* is the mean of the von Mises distribution, i.e., the mean value of *x_m_*. *κ_m_* is a positive number characterizing the concentration of the distribution, and controls the reliability of cue *x_m_*.

The prior *p*(*s*_1_,*s*_2_) describes the probability of concurrence of stimulus features (*s*_1_, *s*_2_) coming from the same stimulus, and it determines the extent to which the two stimulus features should be integrated. In this study, we consider a prior which has been used in several multisensory integration studies^11,20–22^, which is written as

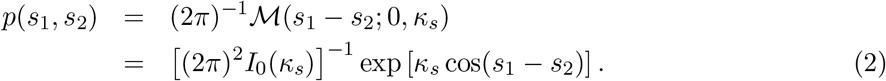

This prior reflects that the two stimulus features from the same stimulus tend to have similar values. The parameter *κ_s_* specifies the concurrence probability of two stimulus features, and determines the extent to which the two cues should be integrated. In the limit *κ_s_* → ∞, it will lead to full integration (see, e.g., ref. 5). Note that the marginal prior *p*(*s_m_*) is a uniform distribution according to the definition.

It has been revealed that the brain integrates visual and vestibular cues to infer heading direction in a manner close to Bayesian inference^8,9^. Following Bayes’ theorem, optimal multisensory integration is achieved by computing the posterior of two stimuli according to

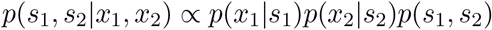

Since the calculations of the two stimuli are exchangeable, hereafter we only present the results for *s*_1_. The posterior of *s*_1_ is calculated through marginalizing the joint posterior in the above equation,

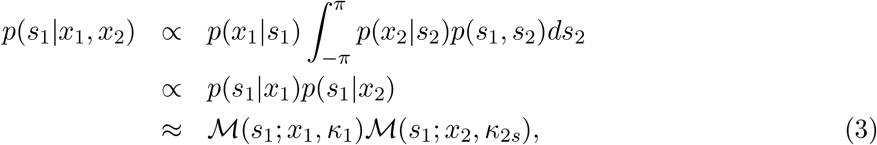

where we have used the conditions that the marginal prior distributions of *s_m_* and *x_m_* are uniform, i.e., *p*(*s_m_*) = *p*(*x_m_*) = (2π)^−1^. Note that 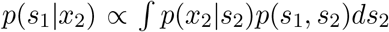 is approximated to be 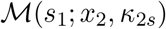 through equating the mean resultant length of distribution (Eq. 12)^23^.

The above equation indicates that in multisensory integration, the posterior of a stimulus given combined cues is equal to the product of the posteriors given the individual cues. Notably, although *x*_1_ and *x*_2_ are generated independently by *s*_1_ and *s*_2_ (since the visual and vestibular signal pathways are separated), *x*_2_ also provides information of *s*_1_ due to the correlation between *s*_1_ and *s*_2_ specified in the prior.

Finally, since the product of two von Mises distributions is again a von Mises distribution, the posterior distribution is 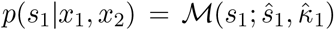, whose mean and concentration can be obtained from its moments given by

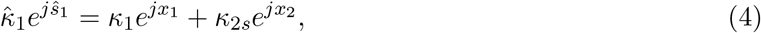

where *j* is an imaginary number. Eq. 4 is the result of Bayesian optimal integration in the form of von Mises distributions, and they are the criteria to judge whether optimal cue integration is achieved in the neural system. A link between the Bayesian criteria for von Mises and Gaussian distributions are presented in SI.

Eq. 4 indicates that the von Mises distribution of a circular variable can be interpreted as a vector in a two-dimensional space with its mean and concentration representing the angle and length of the vector, respectively (Fig. 3A). In this interpretation, the product of two von Mises distributions can be represented by the summation of the corresponding two vectors. Thus, optimal multisensory integration is equivalent to vector summation (see Eq. 4), with each vector representing the posterior of the stimulus given each cue (the sum of the two green vectors yields the blue vector in Fig. 3B).

**Figure 3:**
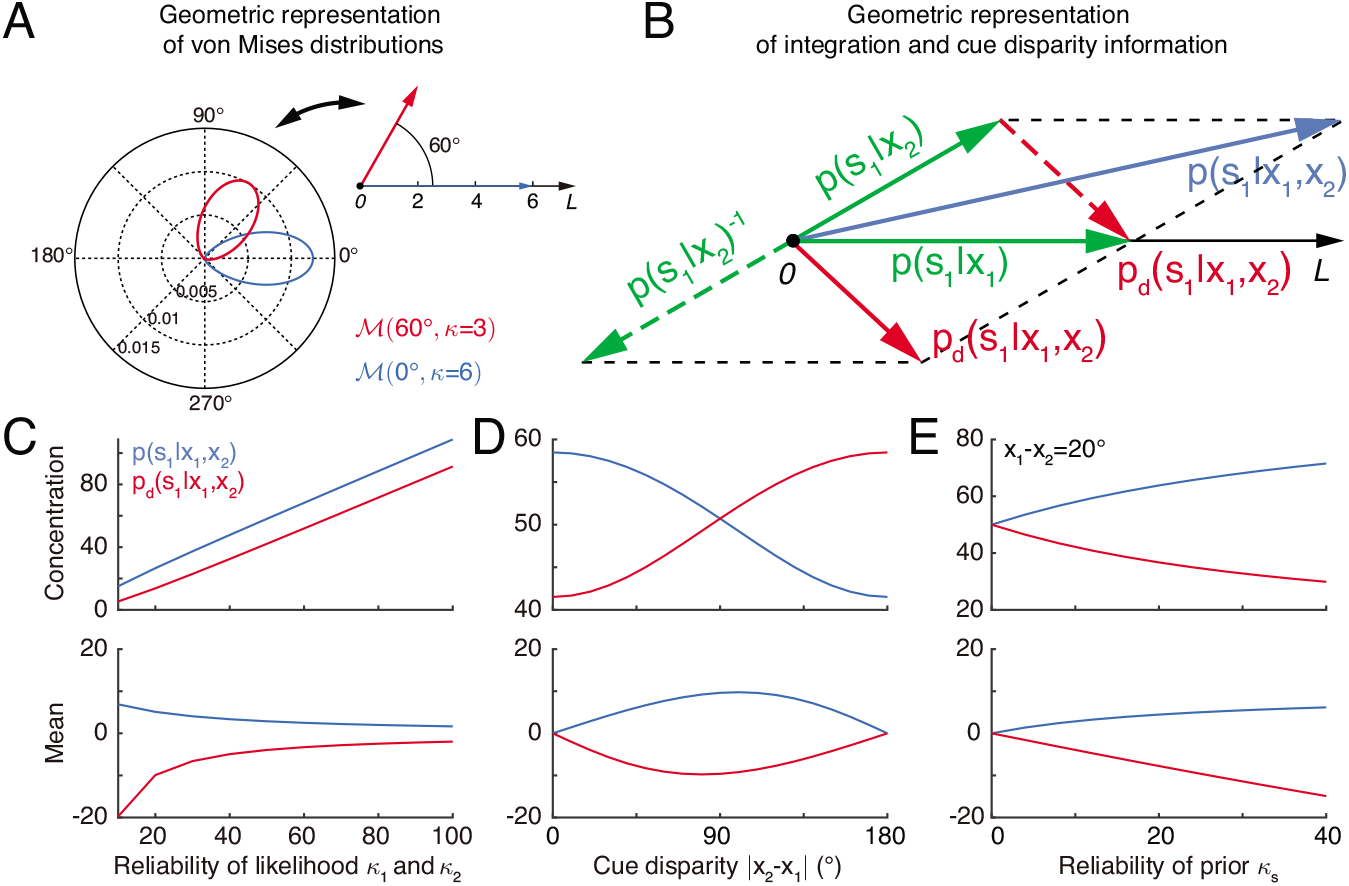
Geometric interpretation of multisensory processing of circular variables. (A) Two von Mises distributions plotted in the polar coordinate (bottom-left) and their corresponding geometric representations (top-right). A von Mises distribution can be represented as a vector, with its mean and concentration corresponding to the angle and length of the vector, respectively. (B) Geometric interpretation of cue integration and the cue disparity information. The posteriors of s1 given single cues are represented by two vectors (green). Cue integration (blue) is the sum of the two vectors (green), and the cue disparity information (red) is the difference of the two vectors. (C-E) The mean and concentration of the integration (blue) and the cue disparity information (red) as a function of the cue reliability (C), cue disparity (D), and reliability of prior (E). In all plots, *κ_s_* = 50, *κ*_1_ = *κ*_2_ = 50, *x*_1_ = 0° and *x*_2_ = 20°, except that the variables are *κ*_1_ = *κ*_2_ in C, *x*_2_ in D, and *κ*_s_ in E.

#### Probabilistic model of multisensory segregation

The above probabilistic model for multisensory integration assumes that sensory cues are originated from the same stimulus. In case they come from different stimuli, the cues need to be segregated, and the neural system needs to infer stimuli based on individual cues. In practice, the brain needs to differentiate these two situations. In order to achieve reliable and rapid multisensory processing, we propose that while integrating sensory cues, the neural system simultaneously extracts the disparity information between cues, so that with this complementary information, the neural system can assess the validity of cue integration.

An accompanying consequence of multisensory integration is that the stimulus information associated with individual cues is lost once they are integrated (see Supplementary Fig. S1). Hence besides assessing the validity of integration, extracting both congruent and disparity information by simultaneous integration and segregation enables the system to recover the lost information of individual cues if needed.

The disparity information of stimulus 1 obtained from the two cues is defined to be

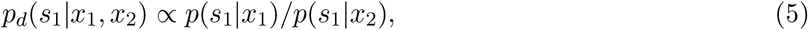

which is the ratio between the posterior given two cues and hence measures the discrepancy between the estimates from different cues. By taking the expectation of log *p_d_* over the distribution *p*(*s*_1_|*x*_1_), it gives rise to the Kullback-Leibler divergence between the two posteriors given each cue. This disparity measure was also used to discriminate alternative moving directions in ref. 24.

Utilizing the property of the von Mises distribution and the periodicity of heading directions (–cos(*s*_1_ – *x*_2_) = cos(*s*_1_ – *x*_2_ – π)), Eq. 5 can be re-written as

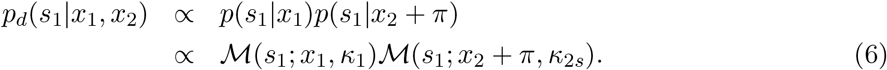

Thus, the disparity information between two cues can also be expressed as the product of the posterior given the direct cue and the posterior given the indirect cue with the cue direction shifted by π. Indeed, analogous to the derivation of Eq. 3, Eq. 6 can be deduced in the same framework as multisensory integration but with the stimulus prior *p*(*s*_1_, *s*_2_) being modified by a shift π in the angular difference. Similarly, 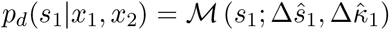 whose mean and concentration can be derived as

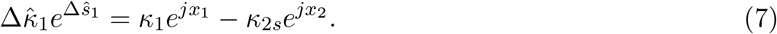

The above equation is the criteria to judge whether the disparity information between two cues is encoded in the neural system.

Similar to the geometrical interpretation of multisensory integration, multisensory segregation is interpreted as vector subtraction (the subtraction between two blue vectors yields the red vector in Fig. 3B). This enables us to assess the validity of multisensory integration. When the two vectors representing the posteriors given the individual cues have small disparity, i.e., the estimates from individual cues tend to support each other, the length of the summed vector is long, implying that the posterior of cue integration has a strong confidence, whereas the length of the subtracted vector is short, implying that the weak confidence of two cues are disparate (Fig. 3D). If the two vectors associated with the individual cues have a large disparity, the interpretation becomes the opposite (Fig. 3D). Thus, by comparing the lengths of the summed and subtracted vectors, the neural system can assess whether two cues should be integrated or segregated.

Figs. 3C and E further describes the integration and segregation behaviors when the model parameters vary. As shown in Fig. 3C, when the likelihoods have weak reliabilities, the Bayesian estimate relies more on the prior. Since the prior encourages integration of the two stimuli, the posterior estimate of stimulus 1 becomes more biased towards cue 2. At the same time, the mean of the disparity information is biased towards the angular difference of the likelihood peaks. On the other hand, when the likelihoods are strong, the Bayesian estimate relies more on the likelihood, and the posterior estimate of stimulus 1 becomes less biased towards cue 2. The behavior when the prior concentration *κ*_s_ varies can be explained analogously (Fig. 3E).

A notable difference between von Mises distribution and Gaussian distribution is that the concentration of integration and disparity information changes with cue disparity in von Mises distribution (Fig. 3D), while they are fixed in Gaussian distribution^25^.

### Neural implementation of cue integration and segregation

Before introducing the neural circuit model, we first describe intuitively how opposite neurons encode the cue disparity information and the motivation of the proposed network structure.

Optimal multisensory integration computes the posterior of a stimulus given combined cues according to Eq. 3, which is equivalent to solving the equation ln*p*(*s*_i_|*x*_i_, *x*_2_) = ln *p*(*s*_i_|*x*_1_) + ln *p*(*s*_i_|*x*_2_). Ma *et al*.found that under the conditions that neurons fire independent Poisson spikes, the optimal integration can be achieved by combining the neuronal resposnes under single cue conditions, that is **r**_*j*_(*x*_i_,*x*_2_) = **r**_*j*_(*x*_1_) +**r**_*j*_(*x*_2_) (see details in SI), where **r**(*x*_1_,*x*_2_) and **r**(*x*_m_) are the responses of a population of neurons to the combined and single cues respectively ^12^. Ma *et al*. further demonstrated that such a response property can be approximately achieved in a biological neural network. Similarly, multisensory segregation computes the disparity information between cues according to ln *p_d_*(*s*_1_|*x*_1_,*x*_2_) = ln *p*(*s*_i_|*x*_1_)+ln *p*(*s*_i_|*x*_2_ +π) (see Eq. 6). Analogous to multisensory integration, the optimal segregation can be achieved by **r**_*j*_(*x*_1_,*x*_2_) = **r**_*j*_(*x*_1_) + **r**_*j*’_(*x_2_*), where the preferred stimulus of neurons satisfying *θ_j’_* = *θ_j_* + π (see details in SI). That is, the neurons combine the responses to the direct cue and the responses to the indirect cue but shifted to opposite direction. This inspires us to consider a network model where the inputs of indirect cue received by opposite neurons are shifted to opposite direction via connections. Below, we present the network model and demonstrate that the opposite neurons emerge from the connectivity and are able to achieve optimal segregation.

#### The decentralized neural network model

The neural circuit model we consider has the decentralized structure^11^, in the sense that it consists of two reciprocally connected modules (local processors), representing MSTd and VIP respectively (Fig. 4A). Each module carries out multisensory processing via cross-talks between modules. This decentralized architecture agrees with the experimental findings that neurons in MSTd and VIP both exhibit multisensory responses and that the two areas are abundantly connected with each other^15,16^. Below we only describe the key features of the decentralized network model, and its detailed mathematical description is presented in Methods (Eqs. 14-20).

**Figure 4:**
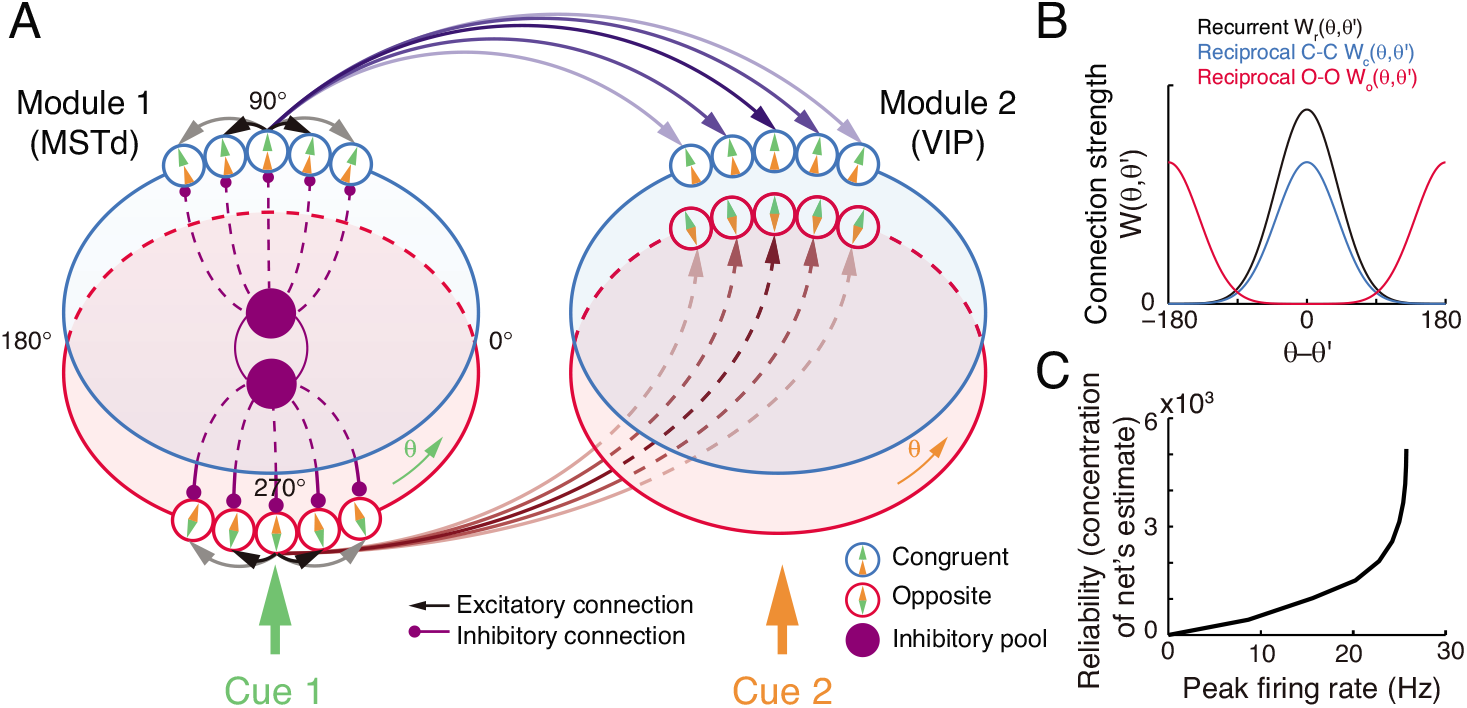
The decentralized neural circuit model for multisensory processing. (A) The network consists of two modules, which can be regarded as MSTd and VIP respectively. Each module has two groups of excitatory neurons, congruent (blue circles) and opposite neurons (red circles). Each group of excitatory neurons are connected recurrently with each other, and they are all connected to an inhibitory neuron pool (purple disk) to form a continuous attractor neural network. Each module receives a direct cue through feedforward inputs. Between modules, congruent neurons are connected in the congruent manner (blue arrows), while opposite neurons are connected in the opposite manner (brown lines). (B) Connection profiles between neurons. Black line is the recurrent connection pattern between neurons of the same type in the same module. Blue and red lines are the reciprocal connection patterns between congruent and opposite neurons across modules respectively. (C) The reliability of the networks estimate of a stimulus is encoded in the peak firing rate of the neuronal population. Typical parameters of network model: *ω* = 3 × 10^−4^, *J_int_* = 0.5, *J_rc_* = 0.3*J_c_*, *J_rp_* = 0.5*J_rc_*, *I_b_* and *F* in Eq. 20 are 1 and 0.5 respectively.

At each module, there exist two groups of excitatory neurons: congruent and opposite neurons (blue and red circles in Fig. 4A respectively), and they have the same number of neurons, as supported by experiments (Fig. 2C)^18,19^. Each group ofneurons is modelled as a continuous attractor neural network (CANN), mimicking the encoding of heading-direction in neural systems^26,27^. In CANN, each neuron is uniquely identified by its preferred heading direction θ with respect to the direct cue conveyed by feedforward inputs. The neurons in the same group are recurrently connected, and the recurrent connection strength between neurons *θ* and *θ’* is modelled as a von Mises function decaying with the disparity between two neurons’s preferred directions |*θ* — *θ’*| (Fig. 4B black line and Eq. 15). In the model, the recurrent connection strength is not very strong to support persistent activities after switching off external stimuli, because no persistent activity is observed in multisensory areas. Moreover, neuronal responses in the same group are normalized by the total activity of the population (Eq. 18), called divisive normalization^28^, mimicking the effect of a pool of inhibitory neurons (purple disks in Fig. 4B). Each group of neurons has its individual inhibitory neuron pool, and the two pools of inhibitory neurons in the same module share their overall activities (Eq. 19), which intends to introduce mutual inhibition between congruent and opposite neurons.

Between modules, neurons of the same type are reciprocally connected with each other (Figs. 4A-B). For congruent neurons, they are connected with each other in congruent manner (Eq.16 and Fig. 4B blue line), that is, the more similar their preferred directions are, the stronger the neuronal connection is. For opposite neurons, they are connected in the opposite manner (Eq. 17 and Fig. 4B red line), that is, the more different their preferred directions are, the stronger the neuronal connection is. Since the maximum difference between two circular variables is π, an opposite neuron in one module preferring *θ* has the strongest connection to the opposite neuron preferring *θ* + π in the other module. This agrees with our intuitive understanding as described above (as suggested by Eq. 6): to calculate the disparity information between two cues, the neuronal response to the combined cues should integrate its responses to the direct cue and its response to the indirect one but with the cue direction shifted by π (through the offset reciprocal connections). We set the connection profile between the opposite neurons to be of the same strength and width as that between the congruent ones (comparing Eqs. 16 and 17), ensuring that the tuning functions of the opposite neurons have the similar shape as those of the congruent ones, as observed in the experimental data^18^.

When sensory cues are applied, the neurons combine the feedforward, recurrent, and reciprocal inputs to update their activities (Eq. 14), and the multisensory integration and segregation will be accomplished by the reciprocal connections between network modules. The results are presented below.

#### Tuning properties of congruent and opposite neurons

Simulating the neural circuit model, we first checked the tuning properties of neurons. The simulation results for an example congruent neuron and an example opposite neuron in module 1 responding to single cues are presented in Fig. 5. It shows that the congruent neuron, in response to either cue 1 or cue 2, prefers the same direction (—90°) (Fig. 5A), whereas the opposite neuron, while preferring —90° for cue 1, prefers 90° for cue 2 (Fig. 5B). Thus, the tuning properties of congruent and opposite neurons naturally emerge through the network dynamics.

**Figure 5:**
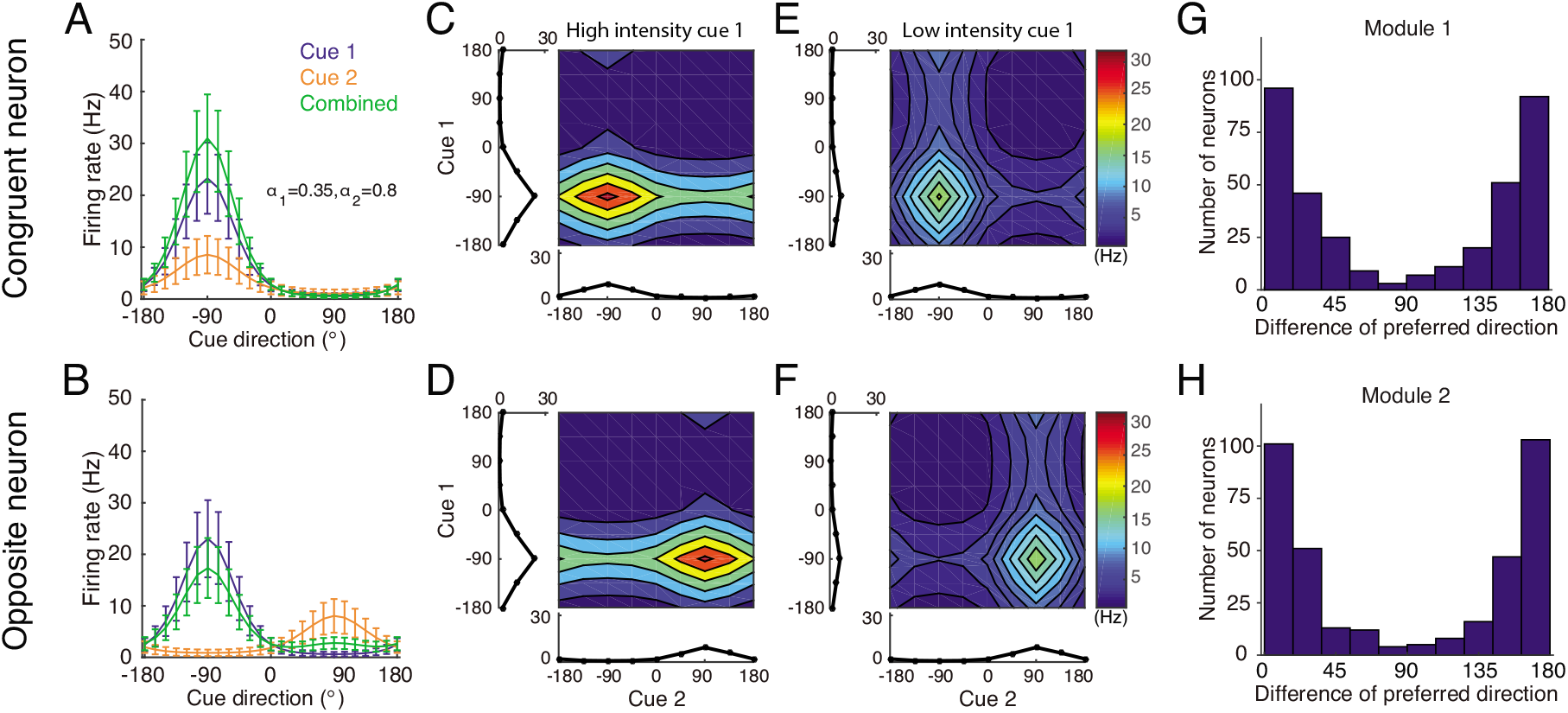
Tuning properties of congruent and opposite neurons in the network model. (A-B) The tuning curves of an example congruent neuron (A) and an example opposite neuron (B) in module 1 under three cueing conditions. (C-D) The bimodal tuning properties of the example congruent (C) and the example opposite (D) neurons when cue 1 has relatively higher reliability than cue 2 in driving neurons in module 1, with α1 = 0.58*α*2, where *αm* is the amplitde of cue *m* given by Eq. 20. The two marginal curves around each contour plot are the unimodal tuning curves. (E-F) Same as (C-D), but cue 1 has a reduced reliability with *α*1 = 0.12*α*2. (G-H) The histogram of the differences of neuronal preferred directions with respect to two cues in module 1 (G) and module 2 (H), when the reciprocal connections across network modules contain random components of roughly the same order as the connections. Parameters: (A-B) *α*1 = 0.35*U*_0_, and *α*2 = 0.8*U*_0_; (C-F) *α*2 = 1.5*U*_0_.*α*1 = 0.35*U*_0_ in (C-D) while *α*1 = 0.1*U*_0_ in (E-F). Other parameters are the same as those in Fig. 4.

We further checked the responses of neurons to combined cues, and found that when there is no disparity between the two cues, the response of a congruent neuron is enhanced compared to the single cue conditions (green line in Fig. 5A), whereas the response of an opposite neuron is suppressed compared to its response to the direct cue (green line in Fig. 5B). These properties agree with the experimental data^8,9^ and is also consistent with the interpretation that the integrated and segregated amplitudes are respectively proportional to the vector sum and difference in Fig. 3. Following the experimental protocol ^13^, we also plotted the bimodal tuning curves of the example neurons in response to the combined cues of varying reliability, and observed that when cue 1 has a relatively high reliability, the bimodal responses of both neurons are dominated by cue 1 (Fig. 5C-D), indicating that the neuronal firing rates are affected more significantly by varying the angle of cue 1 than by that of cue 2, whereas when the reliability of cue 1 is reduced, the result becomes the opposite (Fig. 5E-F). These behaviors agree with the experimental observations ^13^.

Apart from the congruent and opposite neurons, the experiments also found that there exist a portion of neurons, called intermediate neurons, whose preferred directions to different cues are neither exactly the same nor the opposite, but rather have differences in between 0° and 180°^18,19^. We found that by considering the realistic imperfectness of neuronal reciprocal connections (e.g., adding random components in the reciprocal connections in Eqs. (16 and 17), see Methods), our model reproduced the distribution of intermediate neurons as observed in the experiment (Fig. 5G-H)^18,19^.

#### Optimal cue integration and segregation via congruent and opposite neurons

In response to the noisy inputs in a cueing condition, the population activity of the same group of neurons in a module exhibits a bump-shape (Fig. 6A), and the position of the bump is interpreted as the network’s estimate of the stimulus (Fig. 6B)^27,29,30^. In a single instance, we used the populationvector to read out the stimulus value (Eq. 21)^31^. The statistics ofthe bump position sampled from a collection of instances reflects the posterior distribution of the stimulus estimated by the neural population under the given cueing condition. Note that in this probabilistic population coding scheme, the concentration of the decoded posterior distribution is independent of the widths of the bumps at individual instances.

**Figure 6:**
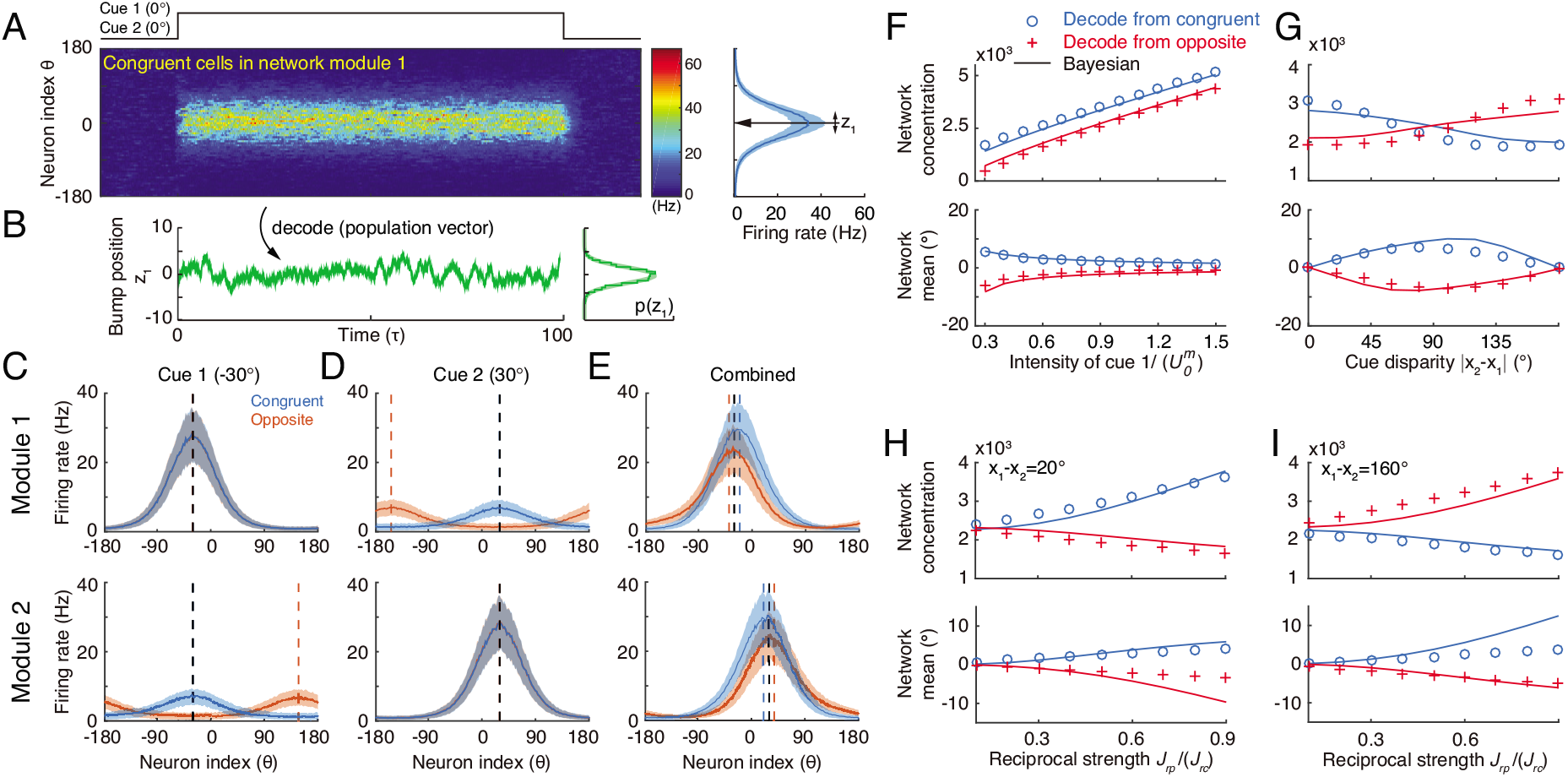
Optimal cue integration and segregation collectively emerge in the neural population activities in the network model. (A) Illustration of the population response of congruent neurons in module 1 when both cues are presented. Color indicates firing rate. Right panel is the temporal average firing rates of the neural population during cue presentation, with shaded region indicating the standard deviation (SD). (B) The position of the population activity bump at each instance is interpreted as the network’s estimate of the stimulus, referred to as *z*1, which is decoded by using population vector. Right panel is the distribution of the decoded network’s estimate during cue presentation. (C-E) The temporal average population activities of congruent (blue) and opposite (red) neurons in module 1 (top row) and module 2 (bottom row) under three cueing conditions: only cue 1 is presented (C), only cue 2 is presented (D), and both cues are simultaneously presented (E). (F-I) Comparing the estimates from congruent and opposite neurons in module 1 with the Bayesian predictions, with varying cue intensity (F), with varying cue disparity (G), and with varying reciprocal connection strength between modules (H&I). Symbols: network results; lines: Bayesian prediction. The Bayesian predictions for the estimates of congruent and opposite neurons are obtained by Eq. 4 and Eq. 7. Parameters: (A-E) *α*_1_ = *α*_2_ = 0.35*U*_0_ ; (F) *α*_2_ = 0.7*U*_0_; (G-I) *α*_1_ = *α*_2_ = 0.7*U*_0_, and others are the same as those in Fig. 4. In (F-H), *x*_1_ = 0°, *x*_2_ = 20° and in (I), *x*_1_ = 0°, *x*_2_ = 160°.

To validate the hypothesis that congruent and opposite neurons are responsible for optimal cue integration and segregation respectively, we carried out simulations following the protocol in multisensory experiments ^1^, that is, we first applied individual cues to the network and decoded the network’s estimate of the stimulus through population vector (see details in Methods). With these results, the Bayesian predictions for optimal integration and segregation were calculated according to Eq. 4 and Eq. 7 respectively; we then applied the combined cues to the network, decoded the network’s estimate, and compared them with the Bayesian predictions.

Let us first look at the network’s estimate under single cue conditions. Consider the case that only cue 1 is presented to module 1 at −30°. The population activities of congruent and opposite neurons at module 1 are similar, both centered at —30° (Fig. 6C top), since both types of neurons receive the same feedforward input. On the other hand, in module 2, congruent neurons’ responses are centered at −30°, while opposite neurons’ responses are centered at 150° due to the offset reciprocal connections (Fig. 6C bottom). Similar population activities exist under cue 2 condition (Fig. 6D).

We further look at the the network’s estimate under the combined cue condition. Consider the case that cues 1 and 2 are simultaneously presented to the network at the directions −30° and 30° respectively.Then the disparity between the two cues is 60°, which is less than 90°. Compared with single cue conditions, the responses of congruent neurons are enhanced (comparing Fig. 6E with 6C-D), reflecting the increased reliability of the estimate after cue integration. Indeed, the decoded distribution from congruent neurons sharpens in the combined cue condition and moves to a location between cue 1 and cue 2 (Fig. S2 green), which is a typical phenomenon associated with cue integration. In contrast, with combined cues, the responses of opposite neurons are suppressed compared with those of the direct cue (comparing Fig. 6E with 6C-D). Certainly, the distribution of cue disparity information decoded from opposite neurons in combined cue condition is wider than that that under the direct cue condition (Fig. S2 purple). Note that when the cue disparity is larger than 90°, the relative response of congruent and opposite neurons will be reversed (results are not shown here).

To demonstrate that the network implements optimal cue integration and segregation and how the network encodes the probabilistic model (Eqs. 1 and 2), we changed a parameter at a time, and then compared the decoded results from congruent and opposite neurons with the Bayesian prediction. Fig.6F-I indicates that the network indeed implements optimal integration and segregation. Moreover, comparing the network results with the results of the probabilistic model, we could find the analogy that the input intensity encodes the reliability of the likelihood (Eq. 1, comparing Fig. 6F with Fig. 3C), and the reciprocal connection strength effectively represents the reliability of the prior (Eq. 2, comparing Fig. 6H with Fig. 3E), which is consistent with a previous study^11^. We further systematically changed the network and input parameters over a large parameter region and compare the network results with Bayesian prediction. Our results indicated that the network model achieves optimal integration and segregation robustly over a large range of parameters (Fig. S3), as long as the connection strengths are not so large that winner-take-all happens in the network model.

### Concurrent multisensory processing

The above results elucidate that congruent neurons integrate cues, whereas opposite neurons compute the disparity between cues. Based on these complementary information, the brain can access the validity of cue integration and can also recover the stimulus information associated with single cues lost due to integration. Below, rather than exploring the detailed neural circuit models, we demonstrate that the brain has resources to implement these two operations based on the activities of congruent and opposite neurons.

#### Assessing integration vs. segregation

The competition between congruent and opposite neurons can determine whether the brain should integrate or segregate two cues. Fig. 7A displays how the mean firing rates of two types of neurons change with the cue disparity, which shows that the activity of congruent neurons decreases with the disparity, whereas the activity of opposite neurons increases with the disparity, and they are equal at the disparity value of 90°. The brain can judge the validity of integration based on the competition between these two groups of neurons (see more remarks in Conclusions and Discussions). Specifically, the group of congruent neurons wins when the cue disparity is small, indicating the choice of integration, and the group of opposite neurons wins when the cue disparity is large, indicating the choice of segregation. The decision boundary is at the disparity of 90°, if the activities of congruent and opposite neurons have equal weights in decision-making. In reality, however, the brain may assign different weights to congruent and opposite neurons and realize a decision boundary at the position satisfying the statistics of inputs (Fig. 7B).

**Figure 7:**
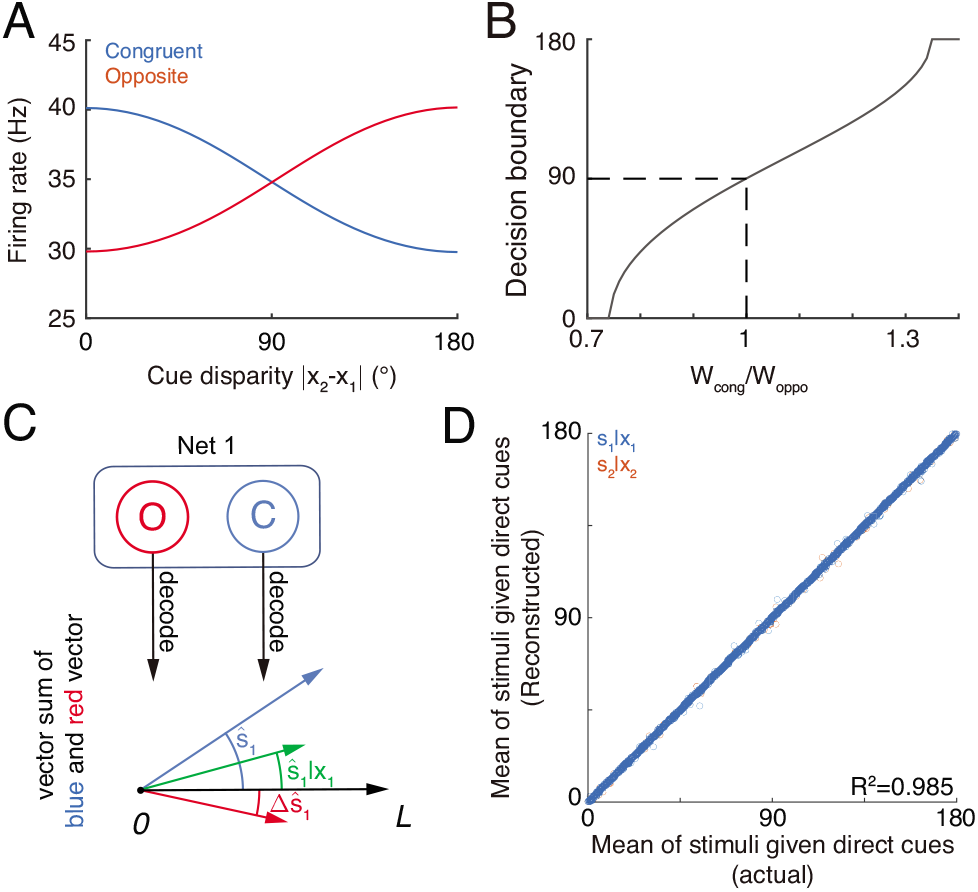
Concurrent multisensory processing with congruent and opposite neurons. (A-B) Accessing integration versus segregation through the joint activity of congruent and opposite neurons. (A) The firing rate of congruent and opposite neurons exhibit complementary changes with cue disparity *x*_1_ - *x*_2_. (B) The decision boundary of the competition between congruent and opposite neurons changes with read out weight from congruent *W_cong_* and opposite neurons *W_oppo_*. It is given by the value of *x*_1_ - *x*_2_ at which 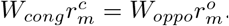. Dashed line is when *W_cong_* = *W_oppo_*, the decision boundary is at 90°. (C-D) Recovering single cue information from two types of neurons. (C) Illustration of recovering through the joint activities of congruent (blue) and opposite (red) neurons under the combined cue condition. We decoded the estimate from congruent and opposite neurons respectively, and then vector sum the decoded results recovering the single cue information. (D) Comparing the recovered mean of the stimulus given the direct cue with the actual value. Parameters: those in (A-B) are the same as those in Fig. 6A, and those in D are the same as those in Fig. S3.

#### Recovering the single cue information

Once the decision for cue segregation is reached, the neural system at each module needs to decode the stimulus based purely on the direct cue, and ignores the irrelevant indirect one. Through combining the complementary information from congruent and opposite neurons, the neural system can recover the stimulus estimates lost in integration, without re-gathering new inputs from lower brain areas if needed (see more remarks in Conclusions and Discussions).

According to Eqs. 3 and 6, the posterior distribution of the stimulus given the direct cue can be recovered by

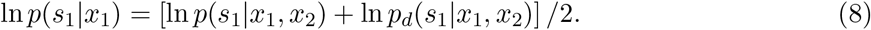

As suggested in refs. 12,24, the above operation can be realized by considering neurons receiving the activities of congruent neurons (representing ln *p*(*s*_1_|*x*_1_, *x*_2_), Fig. 7C blue) and opposite neurons (representing ln*p_d_*(*s*_1_|*x*_1_, *x*_2_), Fig. 7C red) as inputs and generate Poisson spikes, such that the location of population responses and the summed activity encode respectively the mean and variance of the posterior *p*(*s*_1_|*x*_1_) (Fig. 7C green).

Without actually building a neural circuit model, we decoded the stimulus by utilizing the activities of congruent and opposite neurons according to Eq. 8, and compared the recovered result with the estimate of a module when only the direct cue is presented (see the detail in Methods). Fig. 7D further shows that the recovering agrees with actual distribution and is robust against a variety of parameters (*R*^2^ = 0.985). Thus, through combining the activities of congruent and opposite neurons, the neural system can recover the lost stimulus information from direct cues if necessary.

## Conclusions and Discussions

Animals face challenges of processing information fast in order to survive in natural environments, and over millions of years of evolution, the brain has developed efficient strategies to handle these challenges. In multisensory processing, such a challenge is to integrate/segregate multisensory sensory cues rapidly without knowing in advance whether these cues are from the same or different stimuli. To resolve this challenge, we argue that the brain should carry out multisensory processing concurrently by employing congruent and opposite cells to realize complementary functions. Specifically, congruent neurons perform cue integration with opposite neurons computing the cue disparity simultaneously, so that they generate complementary information, based on which the neural system can assess the validity of integration and recover the lost information associated with single cues if necessary. Through this process, the brain can, on one hand, achieve rapid stimulus perception if the cues are from the same stimulus of interest, and on the other hand, differentiate and recognize stimuli based on individual cues with little time delay if the cues are from different stimuli of interest. We built a biologically plausible network model to validate this processing strategy. The model consists of two reciprocally connected modules representing MSTd and VIP, respectively, and it carries out heading-direction inference based on visual and vestibular cues. Our model successfully reproduces the tuning properties of opposite neurons, verifying that opposite neurons encode the disparity information between cues, and demonstrates that the interplay between congruent and opposite neurons can implement concurrent multisensory processing.

Opposite neurons have been found in experiments for years^8,9^, but their functional role remains a mystery. There have been few studies investigating this issue, and two computational works were reported^32,33^, where the authors explored the contribution of opposite neurons in a computational task of inferring self-motion direction by eliminating the confound information of object motion. They showed that opposite neurons are essential, as they provide complementary information to congruent neurons necessary to accomplish the required computation. This result is consistent with our idea that opposite neurons are indispensable in multisensory processing, but our study goes one step further by theoretically proposing that opposite neurons encode the disparity information between cues and that congruent and opposite neurons jointly realize concurrent multisensory processing.

Our hypothesis on the computational role of opposite neurons can be tested in experiments. Through recording the activities of individual congruent neurons in awake monkeys when the monkeys are performing heading-direction discrimination, previous studies demonstrated that congruent neurons implement optimal cue integration^8,9^. We can carry out a similar experiment to check whether opposite neurons encode the cue disparity information. The task is to discriminate whether the disparity from two cues, x**1** – *x*_2_, is either smaller or larger than 0°. To rule out the influence of the change of integrated direction to the activities of neurons, we fix the center of two cues, for example, the center is fixed at 0°, i.e., *x*_1_ + *x*_2_ = 0°, but the disparity between cues *x*_1_ – *x*_2_ varies over trials. Fig. 8A plots the responses of an example opposite neuron and an example congruent neuron respectively in our model with respect to the cue disparity *x*_1_ – *x*_2_. It shows that the firing rate of the opposite neurons changes much more significantly with the cue disparity than that of the congruent neuron, suggesting that the opposite neuron’s response might be more informative to the change of cue disparity compared with a congruent neuron. To quantify how the activity of a single neuron can be used to discriminate the cue disparity, we apply receiver-operating-characteristics (ROC) analysis to construct the neurometric function (Fig. 8B), which measures the fraction of correct discrimination (see Methods). Indeed, the opposite neurons can discriminate the cue disparity much finer than congruent neurons (Fig. 8C). In addition, our model also reproduces the same discrimination task studied in refs. 8,9, i.e., to discriminate whether the heading-direction is on the left or right hand side of a reference direction under different cueing conditions (Fig. S4).

**Figure 8:**
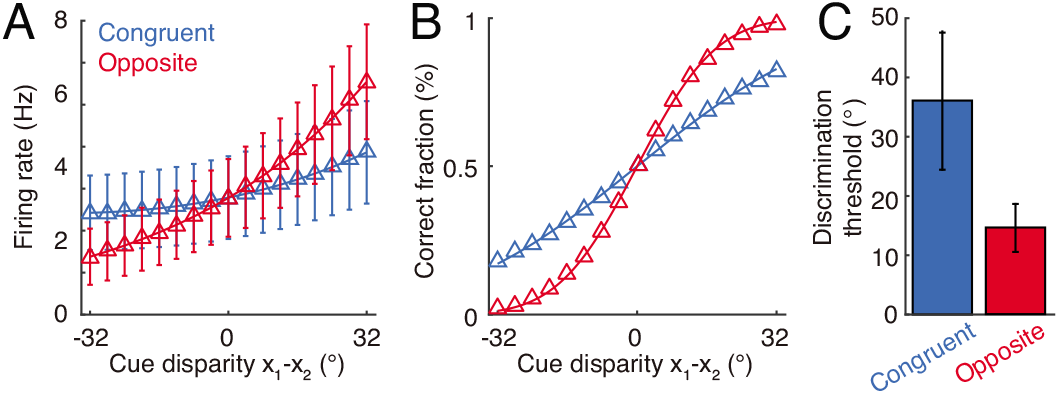
Discrimination of cue disparity by single neurons. (A) The tuning curve of an example congruent (green) and opposite (red) neuron with respect to cue disparity x_1_ – x_2_. In the tuning with respect to cue disparity, the mean of two cues was always at 0°, i.e., x_1_ +x_2_ = 0, while their disparity *x*_1_ – *x*_2_ was varied from −32° to 32° with a step of 4°. The two example neurons are in network module 1, and both prefer 90° with respect to cue 1. However, the congruent neuron prefers 90° of cue 2, while the opposite neuron prefers −90° with respect to cue 2. Error bar indicates the SD of firing rate across trials. (B) The neurometric function of the example congruent and opposite neuron in a discrimination task to determine whether the cue disparity *x*_1_ – *x*_2_ is larger than 0° or not. Lines are the cumulative Gaussian fit of the neurometric function. (C) Averaged neuronal discrimination thresholds of the example congruent and opposite neurons. Parameters: α_1_ = 0.25*U*_0_, α2 = 0.8*U*_0_, and others are the same as those in Fig. 4.

The present study only investigated integration and segregation of two sensory cues, but our model can be generalized to the cases of processing more than two cues that may happen in reality^34^. In such situations, the network model consists of *N* > 2 modules, and in module m, the received sensory cues can be differentiated as the direct one and the integrated results through combining all cues,

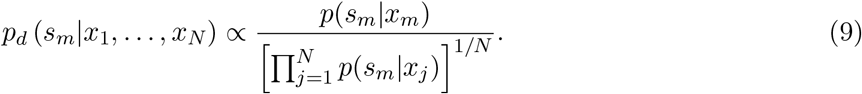

Congruent neurons can be reciprocally connected with each other between modules in the congruent manner as described above, so that they integrate the direct and all indirect cues optimally in the distributed manner. Opposite neurons could receive the direct cue from feedforward inputs (numerator in Eq. 9), and receive the activites of congruent neurons in the opposite manner (denominator in Eq. 9) through offset connection by 180°. The interplay between congruent and opposite neurons determines whether the direct cue should be integrated with all other cues at each module, and their joint activities can recover the stimulus information based only on the direct cue if necessary. This encoding strategy is similar with the norm-based encoding of face found in IT **35** neurons^35^.

In the present study, we only demonstrated by analysis that the neural system can utilize the joint activities of congruent and opposite neurons to assess the validity of cue integration and to recover the information of direct cues in cue integration, but we did not go into the detail of how the brain actually carries out these operations. For assessing the validity of cue integration, essentially it is to compare the activities of congruent and opposite neurons and the winner indicates the choice. This competition process can be implemented easily in neural circuitry. For instance, it can be implemented by considering that congruent and opposite neurons are connected to the same inhibitory neuron pool which induces competition between them, such that only one group of neurons will sustain active responses after competition to represent the choice; alternatively, the activities of congruent and opposite neurons provide competing inputs to a decision-making network, and the latter generates the choice by accumulating evidence over time^36,37^. Both mechanisms are feasible but further experiments are needed to clarify which one is used in practice. For recovering the stimulus information from direct cues by using the activities of congruent and opposite neurons, this study has shown that it can be done in a biologically plausible neural network, since the operation is expressed as solving the linear equation given by Eq. 8. A concern is, however, whether recovering is really needed in practice, since at each module, the neural system may employ an additional group of neurons to retain the stimulus information estimated from the direct cue. An advantage of recovering the lost stimulus information by utilizing congruent and opposite neurons is saving the computational resource, but this needs to be verified by experiments.

The present study focused on investigating the role of opposite neurons in heading-direction inference with visual and vestibular cues as an example. In essence, the contribution of opposite neurons is to retain the disparity information between features to be integrated for the purpose of rapid concurrent processing. We therefore expect that opposite neurons, or their counterparts of similar functions, is a general characteristic of neural information processing where feature integration and segregation are involved. Indeed, it has been found that in the middle temporal cortex (MT), two types of neurons exhibit congruent and opposite tuning properties with respect to moving directions at the center and surrounding of their receptive fields, respectively, and their numbers are comparable^38^. Moreover, MT neurons also exhibit congruent and opposite tunings with respect to binocular disparity and motion parallax, respectively ^39^. We hope that this study gives us insight into understanding the general principle of how the brain integrates/segregates multiple sources of information efficiently and rapidly.

## Methods

### Probabilistic model and its inference

The probabilistic model used in this study is widely adopted in multisensory research^20–22,25^. Suppose that two sensory cues *x*_1_ and *x*_2_ are independently generated by two underlying stimuli *s*_1_ and *s*_2_ respectively. In the example of visual-vestibular cue integration^1^, *s*_1_ and *s*_2_ refer to the underlying visual and vestibular moving direction, while *x*_1_ and *x*_2_ are internal representations of moving direction in the visual and vestibular cortices. Because moving direction is a circular variable, we also assume that both *s_m_* and *x_m_* (*m* = 1, 2) are circular variables distributed in the range (–π, π]. Because each cue is independently generated by the corresponding underlying stimulus, the joint likelihood function can be factorized

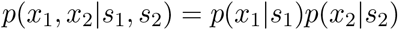

In this study, each likelihood function *p*(*x_m_*|*s_m_*) (*m*= 1,2) is modelled by the von Mises distribution, which is a variant of circular Gaussian distribution^23,40^, given by Eq. 1. Note that in Eq. 1, κm is a positive number characterizing the concentration of the distribution, which is analogous to the inverse of the variance (σ^−2^) of Gaussian distribution. In the limit of large κm, a von Mises distribution 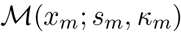 approaches to a Gaussian distribution with variance of 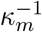^23^. 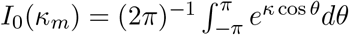 is the modified Bessel function of the first kind and order zero, which acts as the normalization factor of the von Mises distribution.

The prior *p*(*s*_1_,*s*_2_) specifies the probability of occurrence of *s*_1_ and *s*_2_, and is set as a von Mises distribution of the discrepancy between two stimuli^11,20,21^, given by Eq. 2. Note that the marginal prior of either stimulus, e.g., 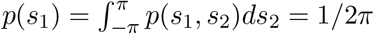 is a uniform distribution.

#### Inference

The inference of underlying stimuli can be conducted by using Bayes’ theorem to derive the posterior

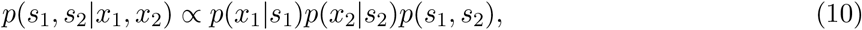

The posterior of either stimuli, e.g., stimulus *s*_1_, can be obtained by marginalizing the joint posterior (Eq. 10) as follows (the posterior of can be similarly obtained by interchanging indices 1 and 2)

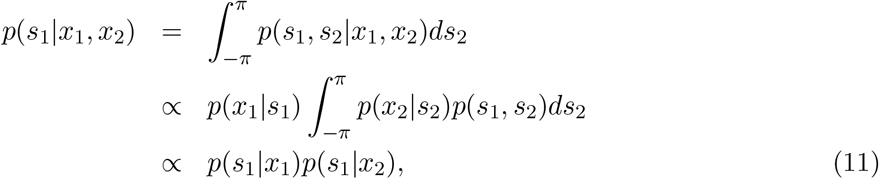

where we used the fact that both marginal distributions *p*(*s_m_*) and *p*(*x_m_*) are uniform and then interchanged the role of *x_m_* and *s*_1_ in their conditional distributions. It indicates that the posterior of *s*_1_ given two cues corresponds to a product of posterior of *s*_1_ when each *x_m_* is individually presented, which could effectively accumulate the information of *s*_1_ from both cues. *p*(*s*_1_|*x*_2_) can be calculated as (see details in SI),

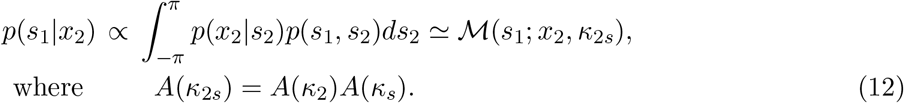

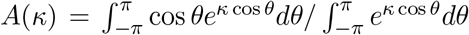 calculates the mean resultant length (first order trigonometric statistics), measuring the dispersion of a von Mises distribution. An approximation was used in the calculation through equating the mean resultant length of the integral with that of a von Mises distribution^23^, because the integral of the product of two von Mises distributions is no longer a von Mises distribution. The meaning of *A*(*κ*_2*s*_) can be understood by considering the Gaussian equivalent of von Mises distribution, where the inverse of concentration *κ*^−1^ can approximate the variance of Gaussian distribution, yielding 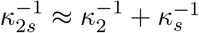.

Finally, substituting the detailed expression into Eq. 11,

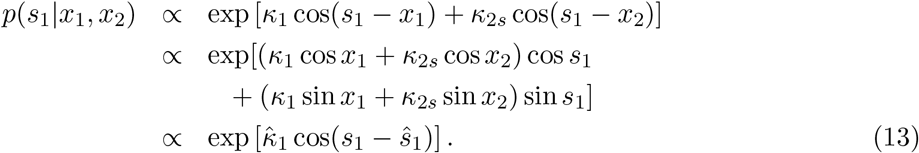

The expressions of the mean *̂S*_1_ and concentration *̂κ*_1_ can be found in Eq. 4. The expressions of △*̂s*_1_ and △*̂κ*_1_ in the disparity information can be similarly calculated and is shown in Eq. 7.

### Dynamics of decentralized network model

We adopted a decentralized network model in this study^11^. The network model contains two network modules, with each module consisting of two groups of neurons with the same number: one is intended to model congruent neurons and another is for opposite neurons. Each neuronal group is modelled as a continuous attractor neural network^27,41,42^, which has been widely used to model the coding of continuous stimuli in the brain^31,43,44^ and it can optimally implement maximal likelihood inference^29,30^. Denote 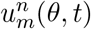 and 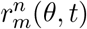 as the synaptic input and firing rate at time t respectively for an *n*-type neuron (*n* = *c*, *o* represents the congruent and opposite neurons respectively) in module *m* (*m* = 1, 2) whose preferred heading direction with respect to the feedforward cue *m* is *θ*. It is worthwhile to emphasize that **θ** is the preferred direction only to the feedforward cue, e.g., the feedforward cue to network module 1 is cue 1, but *θ* does not refer to the preferred direction given another cue, because the preferred direction of an opposite neuron given each cue is different. In the network model, the network module *m* = 1, 2 can be regarded as the brain areas MSTd and VIP respectively. For simplicity, we assume that the two network modules are symmetric, and only present the dynamical equations for network module 1. The dynamical equations for network module 2 can be obtained by interchanging the indices 1 and 2 in the following dynamical equations.

The dynamics of the synaptic input of *n*-type neurons in network module *m*, 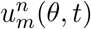, is governed by

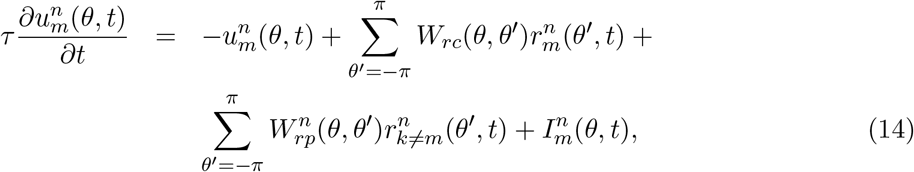

where 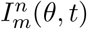 is the feedforward inputs from unisensory brain areas conveying cue information. *W_rc_*(*θ*, *θ*’) is the recurrent connections from neuron *θ*’ to neuron *θ* within the same group of neurons and in the same network module, which is set to be

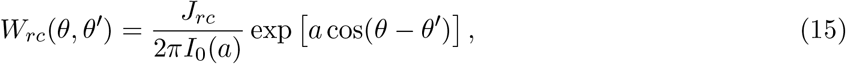

where *a* is the connection width and effectively controls the width of neuronal tuning curves. 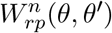 denotes the reciprocal connections between congruent neurons across network modules (*n* = *c*), or between opposite neurons across network modules (*n* = *o*). 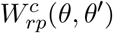 is the reciprocal connections between congruent cells across two modules (the superscript *c* denotes the connections are in a congruent manner, i.e., a 0° neuron will have the strongest connection with a 0° neuron),

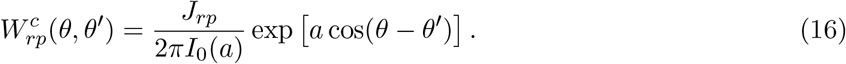

For simplicity, 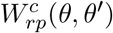 and 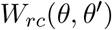 have the same connection width *a*. This simplification does not change the basic conclusion substantially. A previous study indicates that the reciprocal connection strength *J_rp_* determines the extent of cue integration, and effectively represents the correlation of two underlying stimuli in the prior *p*(*s*_1_, *s*_2_)^11^. Moreover, the opposite neurons from different network modules are connected in an opposite manner with an offset of π,

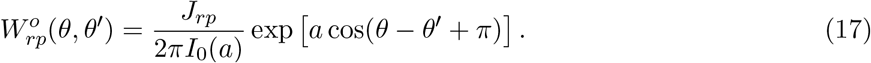

Hence, an opposite neurons preferring 0° of cue 1 in network module 1 will have the strongest connection with the opposite neurons preferring of 180° of cue 2 in network module 2. It is worthwhile to note that the strength and width of 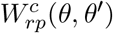 and 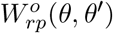 are the same, in order to convey the same information from the indirect cue. This is also supported by the fact that the tuning curves of the congruent and opposite neurons have similar tuning strengths and widths^18^.

Each neuronal group contains an inhibitory neuron pool which sums all excitatory neurons’ activities and then divisively normalize the response of the excitatory neurons,

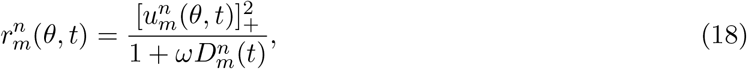

where *ω* controls the magnitude of divisive normalization, and [*x*]_+_ = max(*x*, 0) is the negative rectified function. 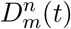 denotes the response of the inhibitory neuron pool associated with neurons of type n in network module *m* at time *t*, which sums up the synaptic inputs of the same type of excitatory neurons 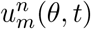 and also receives the inputs from the other type of neurons 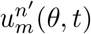,

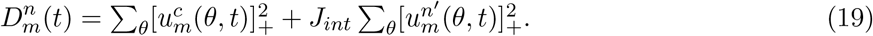

*J_int_* is a positive coefficient not larger than 1, which effectively controls the sharing between the inhibitory neuron pool associated with the congruent and opposite neurons in the same network module. The partial share of the two inhibitory neuron pools inside the same network module introduces competition between two types of neurons, improving the robustness of network.

The feedforward inputs convey the direct cue information from the unisensory brain area to a network module, e.g., the feedforward inputs received by MSTd neurons is from MT which extracts the heading direction from optic flow,

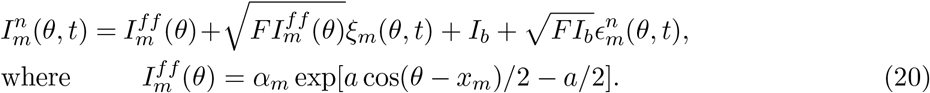

The feedforward inputs contain two parts: one conveys the cue information (the first two terms in above equation), and another the background inputs (the last two terms in the above equation) which are always present no matter whether a cue is presented or not. The variance of the noise in the feedforward inputs 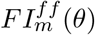 is proportional to their mean, and *F* characterizes the Fano factor. The multiplicative noise is in accordance with the Poisson variability of the cortical neurons’ response. *α_m_* is the intensity of the feedforward input and effectively controls the reliability of cue *m*. *x_m_* is the direction of cue *m*. *I_b_* is the mean of background input.*ξ_m_*(*θ, t*) and 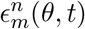 are mutually independent Gaussian white noises of zero mean with variances satisfying 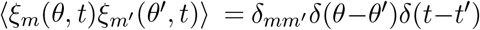 and 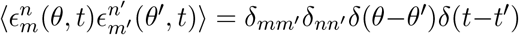 Note that the cue-associated noise *ξm*(*θ, t*) to congruent and opposite neurons are exactly the same, while the background noise 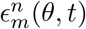 to congruent and opposite neurons are independent of each other. Previous works indicated that the exact form of the feedforward inputs is not crucial, as long as they have a uni-modal shape^42^.

### Network simulation and parameters

Each network module contains 180 congruent and opposite neurons respectively, whose preferred direction with respect to the feedforward cue is uniformly distributed in the feature space (−180°, 180°]. For simplicity, the parameters of the two network modules were chosen symmetric with each other, i.e., all structural parameters of the two modules have the same value. The synaptic time constant τ was rescaled to 1 as a dimensionless number and the time step size was 0.01*τ* in simulation. All connections have the same width *a* = 3, which is equivalent to a value of about 40° for the width of tuning curves of the neurons. The dynamical equations are solved by using Euler method.

The range of parameters was listed in the following if not mentioned otherwise. The detailed parameters for each figure can be found in figure captions. The strength of divisive normalization was *ω* = 3×10^−4^, and *J_int_* = 0.5 which controls the proportion of share between the inhibition pools affiliated with congruent and opposite neurons in the same module (Eq. 19). The absolute values of *ω* and *J_int_* did not affect our basic results substantially, and they only determine the maximal firing rate the neurons can reach. Of the particular values we chose, the firing rate of the neurons saturates at around 50Hz. The recurrent connection strength between neurons ofthe same type and in the same network module was *J_rc_* = [0.3, 0.4]*J*_c_, where *J_c_* is the minimal recurrent strength for a network module to hold persistent activity after switching off feedforward inputs. The expression of *J_c_* can be found in SI. The strength of the reciprocal connections between the network modules is *J_rp_* = [0.1, 0.9]*J_rc_*, and is always smaller than the recurrent connection strength within the same network module. The sum of the recurrent strength *J_rc_* and reciprocal strength *J_rp_* cannot be too large, since otherwise the congruent and opposite neurons in the same network module will have strong competition resulting in the emergence of winner-take-all behavior. However, the winnertake-all behavior was not observed in experiments. The input intensity α was scaled relative to 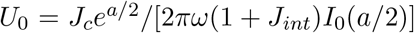, and is distributed in [0.3, 1.5]*U*_0_, where *U*_0_ is the value of the synaptic bump height that a group of neurons can hold without receiving feedforward input and reciprocal inputs when *J_rc_* = *J_c_*. The range of the input intensity was chosen to be wide enough to cover the super-linear to nearly saturated regions of the input-firing rate curve of the neurons. The strength of the background input was *I_b_* = 1, and the Fano factors of feedforward and background inputs were set to 0.5, which led to the Fano factor of single neuron responses taking values of the order 1. In simulations, the position of the population activity bump was read out by calculating the population vector^31,45^. For example, the position of the population activities of the congruent neurons in module 1 at time t was estimated as

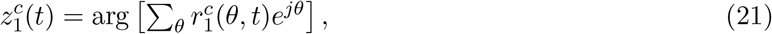

### Demo tasks of network model

#### Testing network’s performance of integration and segregation

We firstly applied each single cue to the network model individually. Under each cueing condition, we recorded the population activities in equilibrium state across time during cue presentation. In equilibrium state, the statistics of neuronal activities across time is equivalent to across trial. For each group of neurons in a module, e.g., the congruent neurons in network module 1, the instantaneous firing activities at an instance are fed into the population vector decoder (Eq. 21) to get the instantaneous stimulus estimate *z*_1_ made by neurons. When the direct cue (cue 1) is presented, the estimates *z*_1_ of a collection of instances are then substituted into Eqs. (S59) and (S61) to calculate the mean and concentration of the activities of the congruent neurons. In the single cue condition, the mean angle of the bump position is effectively the same as *x*_1_. Hence this decoded mean and concentration can be substituted into the first term on the right hand side of Eq. 4. Similarly, when only the indirect cue (cue 2) is presented, the estimates of a collection of instances of neural activities contribute to the second term. The sum of the two terms yields the Bayesian prediction of the optimal integration in combined cue condition. For opposite neurons, we substituted the decoded means and concentrations into Eq. (7) to get the prediction of optimal segregation in combined cue conditon.

#### Reconstructing stimulus estimate under direct cue from congruent and opposite neurons’ activity

The stimulus estimate from its direct cue can be recovered from the joint activities of congruent and opposite neurons in real-time when two cues are simultaneously presented. Eq. 8 indicates that the reconstruction of the posterior distribution of the direct cue can be achieved by multiplying the decoded distribution from congruent and opposite neurons in a network module. Thus, for example, the reconstructed estimate of stimulus 1 at time *t* given its direct cue can be obtained by

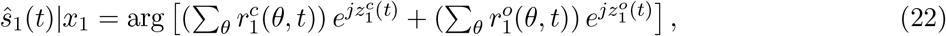

where 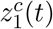 and 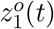 are the positions of the population activities of the congruent and opposite neurons in network module 1 respectively, which were decoded by using population vector (Eq. 21). In real-time reconstruction, the sum of firing rate represents the concentration of the distribution. This is supported by the finding that the reliability of the distribution is encoded by the summed firing rate in probabilistic population code^11,12^.

#### Discriminating cue disparity on single neurons

A discrimination task was designed on the responses of single neurons to demonstrate that opposite neurons encode cue disparity information. The task is to discriminate whether the cue disparity, *x*_1_ — *x*_2_, is either smaller or larger than 0°. In the discrimination task, the mean direction of two cues, *x*_1_ + *x*_2_ = 0, is fixed at 0°, in order to rule out the influence of the change of integrated direction to neuronal activity. Meanwhile, the disparity between two cues, *x*_1_ – *x*_2_, is changed from —32° to 32° with a step of 4°. For each combination of cue direction, we applied three cueing conditions (cue 1, cue 2, combined cues) to the network model for 30 trials and the firing rate distributions of the single neurons were obtained (Fig. 8A and B).

We chose an example congruent neuron preferring 90° in network module 1, and also an example opposite neuron in network module 1 preferring 90° with respect to cue 1. We used receiver operating characteristic (ROC) analysis^46^ to compute the discriminating ability of the example neurons on cue disparity. The ROC value counts the proportion of instances where the direction of cue 1, *x*_1_, is larger than the one of cue 2. Neurometric functions (Fig. 8B and E) were constructed from those ROC values and were fitted with cumulative Gaussian functions by least square, and then the standard deviation of the cumulative Gaussian function was interpreted as the neuronal discrimination threshold (Fig. 8C) ^8^. A smaller value of the discrimination threshold means that the neuron is more sensitive in the discrimination task. Although we adopted the von Mises distribution in the probabilistic model, the firing rate distribution of single neurons can be well fitted by a Gaussian distribution, justifying the use of the cumulative Gaussian distribution to fit the ROC values.

## Acknowledgement

This work is supported by Research Grants Council of Hong Kong (N_HKUST606/12, 605813, 16322616, and 16306817), National Basic Research Program of China (2014CB846101), Natural Science Foundation of China (31261160495), NSF CISE1320651 and IARPA contract D16PC00007.

## Author Contributions

W.H.Z, K.Y.M.W and S.W. designed research; W.H.Z., H.W., K.Y.M.W. and S.W. performed research; A.C., Y.G., and T.S.L. analyzed data and results; W.H.Z., H.W., K.Y.M.W. and S.W. wrote the paper.

